# Activation of innate immune cGAS-STING pathway contributes to Alzheimer’s pathogenesis in 5×FAD mice

**DOI:** 10.1101/2022.10.30.514314

**Authors:** Xiaochun Xie, Guanqin Ma, Xiaohong Li, Jiebin Zhao, Zhen Zhao, Jianxiong Zeng

## Abstract

cGAS senses microbial and host-derived double-stranded DNA (dsDNA) in cytoplasm to trigger cellular innate immune response in a STING-dependent manner. However, it remains unknown whether the cGAS-STING pathway in innate immunity contributes to Alzheimer’s disease (AD). Here we demonstrated the detectable binding of the cGAS-dsDNA in cytoplasm and the activation of microglial cGAS-STING pathway in brains of human AD and aged mice, suggesting a role for the cGAS-STING pathway in this neurodegenerative disease. *Cgas*^−/−^;5×FAD mice were largely protected from cognitive impairment, amyloid-β (Aβ) pathology, neuroinflammation and other sequelae associated with AD. Furthermore, *Cgas* deficiency in microglia inhibited the conversion of astrocytes into neurotoxic A1 phenotype, and thus alleviated oligomeric Aβ peptides-induced neuronal toxicity. Finally, administration of STING inhibitor H-151 potently suppressed the activation of the cGAS-STING pathway and ameliorated AD pathogenesis in 5×FAD mouse model. In conclusion, our present study has identified a critical molecular link between innate immunity and AD, and suggests that therapeutic targeting of the cGAS-STING pathway activity might effectively interfere with the progression of AD.

## Main

Detecting foreign DNA is a critical process of immunity in many organisms. In mammalian cells, this process is mediated mainly by the cyclic GMP-AMP synthase (cGAS)-stimulator of interferon genes (STING) pathway, the mechanistic coupling between the sensing of DNA and the triggering of potent innate immune defense programs^1^. Within this pathway, cGAS binds to double-stranded DNA (dsDNA) allosterically activating its catalytic activity and produces 2′3′ cyclic GMP-AMP (2′3′-cGAMP), a second messenger molecule and potent agonist of STING^2-6^. Activated STING recruits TANK-binding kinase 1 (TBK1) that can phosphorylate STING and the transcription factor interferon regulatory factor 3 (IRF3). Phosphorylation of IRF3 leads to its dimerization and translocation into the nucleus, where phosphorylated IRF3 and nuclear factor κB (NF-κB), the latter being a transcription factor and also activated by STING, function together to initiate the expression of type I interferons (IFNs) and inflammatory cytokines, triggering a broad immune response^1,7^. Nevertheless, the cGAS-STING pathway plays a biological role beyond antimicrobial defense, because cGAS senses and binds to dsDNA in a sequence-independent manner^8^. cGAS can be activated by cytosolic self-DNA derived from genomic or mitochondrial DNA (mtDNA)^1,9^. Accumulated evidence have revealed an expanding role of the cGAS-STING pathway in multiple physiological and pathological processes, including host defense against microbial infections^10^, anti-tumor immunity^11,12^, cellular senescence^13-15^, autophagy^16^, and autoimmune and inflammatory diseases^17^. Recent studies have demonstrated an involvement of the cGAS-STING pathway in several neurological disorders such as ischemic brain injury^18^, Parkinson disease^19^, Huntington disease^20^, and amyotrophic lateral sclerosis^21^. In these disorders, acute or chronic neuroinflammatory response induced by the cGAS-STING pathway contributes to pathological progression of the diseases^22,23^. In particularly, microglial cGAS-STING pathway is likely to play an important role in brain inflammation in tau pathology of Alzheimer’s disease (AD)^24^, the most prevalent neurodegenerative disease. However, it remains unknown whether and how innate immune cGAS-STING pathway participates in the development and progression of amyloid pathology in AD.

AD is characterized by the accumulation of amyloid-β (Aβ) in plaques, aggregation of hyperphosphorylated tau in neurofibrillary tangles and neuroinflammation, together contributing to neurodegeneration and cognitive decline^25^. Neurodegenerative proteins like Aβ can cause oxidative damage of mitochondrial DNA and DNA double-strand breaks (DSBs) in neuronal cultures^26,27^, which potentially increases the level of cytoplasmic dsDNA^28^. cGAS and STING are expressed mainly in microglia, the resident immune cells of the central nervous system (CNS), and thus type I IFNs and inflammatory responses induced by the cGAS-STING pathway are potent in microglia, but less in neurons and astrocytes in the condition of viral infection^29,30^. In this study, we have detected the cytosolic binding of cGAS to dsDNA and the subsequent activation of the cGAS-STING pathway in brains of human AD and aged mice. Either genetic *Cgas* deficiency or administration of STING inhibitor consistently alleviated AD pathogenesis in 5×FAD mouse model, which could be partially attributed to the suppression of neurotoxic A1 astrocytes. These findings are in line with the notion that innate immunity plays a critical role in AD pathogenesis and progression^31^. Thus, our present study has revealed a previously unknown role of the cGAS-STING pathway in amyloid pathology and cognitive impairment in AD.

### The cGAS-STING pathway is activated in brain during AD and aging

Neurodegenerative Aβ can cause oxidative damage of mitochondrial DNA and cellular DSBs^26,27^, which is likely to result in the release of mtDNA fragments into the cytosol. To confirm this, we isolated cytosolic contents of brain cells in 2- or 6-month-old 5×FAD mice, the latter at the stage of abundant Aβ deposition^32^. 5×FAD mouse brain cells had a comparable cytosolic mtDNA (cmtDNA) levels at 2 months old (Extended Data Fig. 1a) but had much higher levels of cmtDNA compared to brain cells from wild-type (WT) mice at 6 months old (Fig. 1a), despite the levels of total mtDNA were very similar (Extended Data Fig. 1b). To examine whether cGAS might bind to cmtDNA in the cytosol and, if any, DNA in the nucleus^33^, we performed a proximity ligation assay (PLA) on brain tissues from 6-month-old 5×FAD mice and detected more than 4-fold of increase in cGAS-dsDNA interactions, as indicated by PLA dots in both cytosol and nucleus of hippocampal region, when compared with WT littermates (Fig. 1b, c). More importantly, we observed similarly cGAS-dsDNA binding events in hippocampal region of human AD samples relative to health control (Extended Data Fig. 1c, d).

**Fig. 1:**
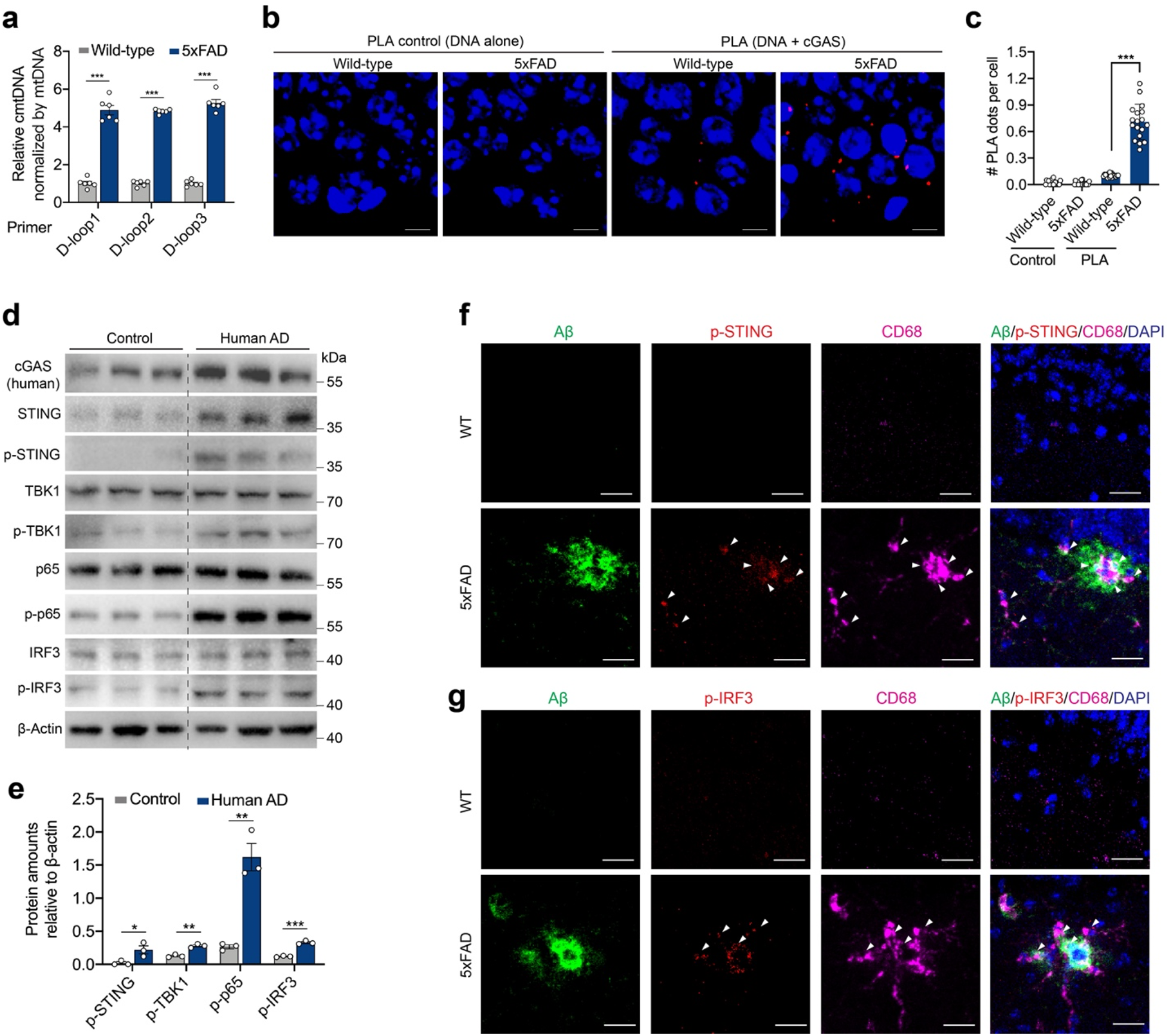
cGAS-STING pathway is activated in brains during AD and aging. **a**, Quantification of cytosolic mitochondrial DNA (cmtDNA) by quantitative PCR (qPCR) in brain cells of 6-month-old 5×FAD mice. Age-matched WT mice were used as control. D-loop indicated a specific fragment within mtDNA. Mean ± SD; *n* = 6; **, *p* < 0.01, Student’s *t*-test. **b**, Proximity ligation assay (PLA) with anti-cGAS and anti-dsDNA antibodies showing the association of cGAS protein with cytosolic or nucleus DNA in hippocampal tissues of 6-month-old 5×FAD mice. Anti-dsDNA antibody alone was used as the PLA control. Age-matched WT mice were used as animal control. Scale bar, 10 μm. **c**, Quantification of PLA red dots per cell in **b**. Three sections in each animal were used to count the dots. Mean ± SD; *n* = 6; *n*.*s*., not significant; ***, *p* < 0.001, one-way ANOVA with Bonferroni’s *post hoc* test. **d**, Western immunoblotting analysis of the expression of the indicated proteins involved in cGAS-SITNG pathway of cortical tissues in human AD. *n* = 3. **e**, Quantification of the expression of the phosphorylated STING (p-STING), TBK1 (p-TBK1), p65 (p-p65), and IRF3 (p-IRF3) relative to β-actin in **d**. Mean ± SD; *n* = 3; *, *p* < 0.05, **, *p* < 0.01, ***, *p* < 0.001, Student’s *t*-test. **f**,**g**, Immunostaining of p-STING (**f**) or p-IRF3 (**g**), as shown as white arrow, with Aβ and activated microglial marker CD68 in DG of 6-month-old WT and 5xFAD mice. Scale bar, 10 μm.

Aging is a primary risk factor for multiple neurodegenerative disorders including AD. We found that brain cells of 20-month-old WT mice had higher levels of cmtDNA relative to their 3-month-old counterparts (Extended Data Fig. 1e), despite having similar levels of total mtDNA (Extended Data Fig. 1f). cGAS-dsDNA binding events were readily detected in hippocampal region of aged mice by PLA assays (Extended Data Fig. 1g, h). Upon sensing dsDNA in the cytosol, cGAS-STING pathway is activated via a series of specific biochemical cascades^1,7^. Immunoblotting assay showed that phosphorylation levels of STING (p-STING), TBK1 (p-TBK1), p65 (p-p65) and IRF3 (p-IRF3) were significantly enhanced in prefrontal cortex of human AD (Fig. 1d, e), as well as in cortex of aged mice (Extended Data Fig. 1i, j). In addition, the downstream type I IFN response in turn induced the expressions of cGAS and STING in both human AD and age mice samples (Fig. 1d and Extended Data Fig. 1i). Importantly, immunostaining assays in brain tissues showed the presence of p-STING (Fig. 1f) and p-IRF3 (Fig. 1g) that was mostly co-localized with activated microglia marker CD68 around Aβ plaques in 5×FAD mice, suggesting the STING-IFN response mainly in microglia. These results demonstrated the binding of cGAS to cytosolic dsDNA and the activation of the cGAS-SITNG pathway in brains during AD and aging.

### *Cgas* deficiency alleviates cognitive impairment and Aβ pathology in 5×FAD mice

To examine the role of the cGAS-STING pathway in AD pathogenesis, we genetically crossed *Cgas*^−/−^ mice with 5×FAD mice, a model for familial AD. 5×FAD mice express human amyloid precursor protein (APP) and presenilin 1 (PSEN1) transgenes with a total of five AD-linked mutations, including the Swedish (K670N/m671l), Florida (I716V), and London (V717I) mutations in APP, and the M146L and L286V mutations in PSEN1. Immunostaining showed that there is no significant difference regarding Aβ burden in 2∼2.5 months old *Cgas*^−/−^;5×FAD mice relative to their *Cgas*^+/+^;5×FAD littermates (Extended Data Fig. 2a, b). However, behavioral tests including nest construction and chow burrowing showed that *Cgas* deficiency improved cognitive functions in 6-month-old 5×FAD mice (Fig. 2a and Extended Data Fig. 2c). We next checked Aβ deposits by measuring Aβ levels using enzyme-linked immunosorbent assay (ELISA). The results revealed that Aβ40 levels in TBS fraction, and Aβ42 levels in both TBS and guanidine fractions were markedly decreased in cortical tissues of *Cgas*^−/−^;5×FAD mice compared to their *Cgas*^+/+^;5×FAD littermates, while Aβ40 levels in guanidine fraction were largely unchanged (Fig. 2b-e). Furthermore, staining with thioflavin S showed a considerable decrease of amyloid cores in hippocampal dentate gyrus (DG) of *Cgas*^−/−^;5×FAD mice compared to their *Cgas*^+/+^;5×FAD littermates (Fig. 2f, g). Additional immunostaining with anti-Aβ antibody revealed the detectable reduction of Aβ deposits in DG of *Cgas*^− /−^;5×FAD mice relative to their littermates (Fig. 2h and Extended Data Fig. 2d, e). Notably, the amount of plaque-associated microglia was much higher in DG of *Cgas*^−/−^;5×FAD mice than that of *Cgas*^+/+^;5×FAD littermates (Fig. 2h, i), suggesting that *Cgas* deficiency improved microglial Aβ clearance.

**Fig. 2:**
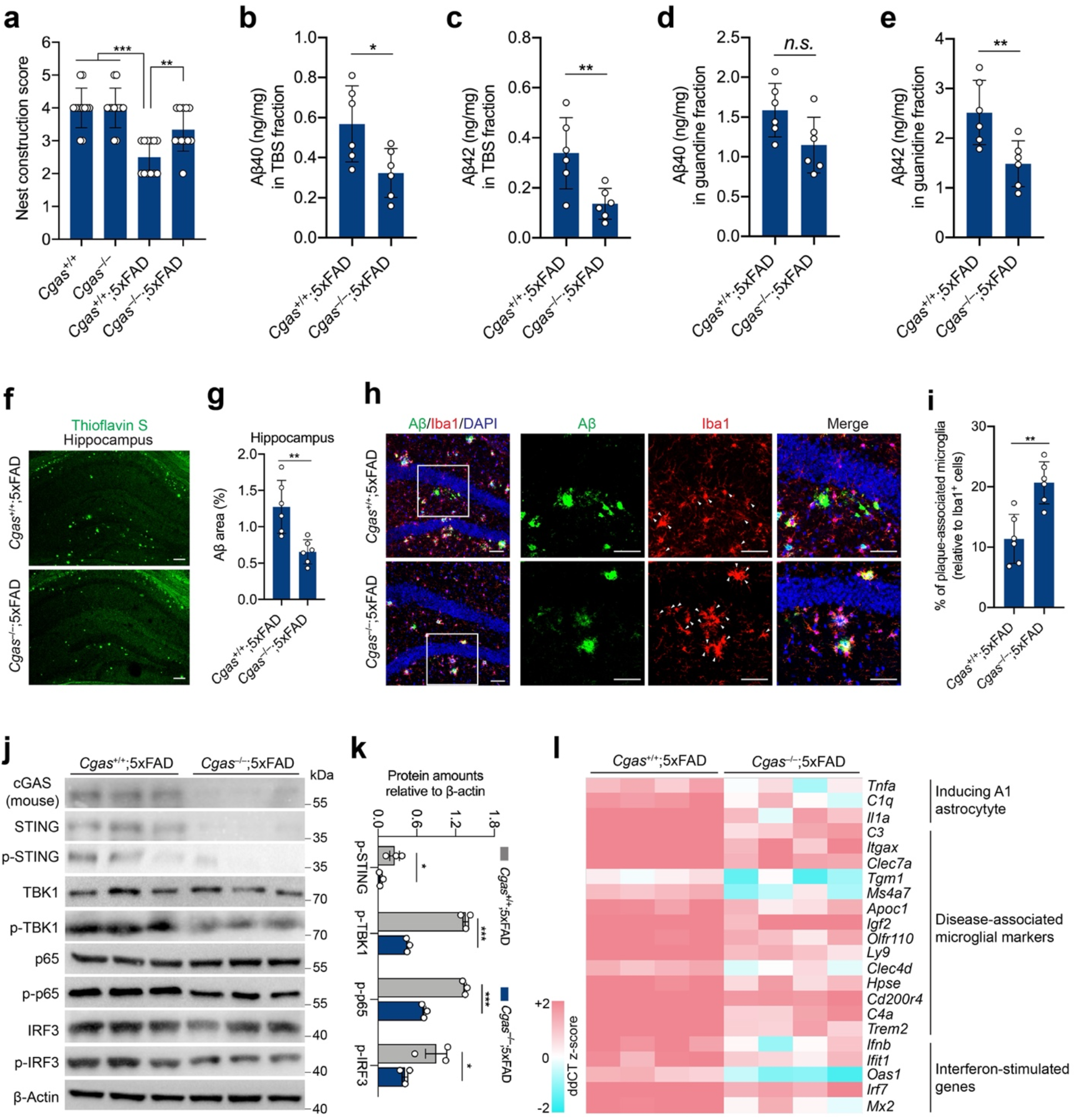
cGAS deficiency alleviates cognitive decline, Aβ pathology and neuroinflammation in 5×FAD mice. **a**, Nest construction score for 6-month-old *Cgas*^+/+^;5×FAD and *Cgas*^−/−^;5×FAD mice. Age-matched *Cgas* ^+/+^ and *Cgas* ^−/−^ mice were used as control. Mean ± SD; *n* = 12; **, *p* < 0.01, ***, *p* < 0.001, one-way ANOVA with Bonferroni’s *post hoc* test. **b-e**, Quantification of the indicated Aβ40 (**b**) and Aβ42 (**c**) in TBS fraction, and Aβ40 (**d**) and Aβ42 (**e**) in guanidine fraction of cortical tissues in 6-month-old *Cgas*^+/+^;5×FAD and *Cgas*^−/−^;5×FAD mice by ELISA. Mean ± SD; *n* = 6; *n*.*s*., not significant; *, *p* < 0.05, **, *p* < 0.01, Student’s *t*-test. **f**, Thioflavin S staining of hippocampal tissues in 6-month-old *Cgas*^+/+^;5×FAD and *Cgas*^−/−^;5×FAD mice. Scale bar, 50 μm. **g**, Quantification of Thioflavin S-labeled amyloid core numbers per mm^2^ in **f**. Mean ± SD; *n* = 6; ***, *p* < 0.001, Student’s *t*-test. **h**, Immunostaining of Aβ and Iba1 in hippocampal dentate gyrus (DG) of 6-month-old *Cgas*^+/+^;5×FAD and *Cgas*^−/−^;5×FAD mice. Scale bar, 40 μm. **i**, Quantification of Aβ plaque-associated Iba1^+^ microglia (indicated by white arrow) in **h**. Mean ± SD; *n* = 6; **, *p* < 0.01, Student’s *t*-test. **j**, Western immunoblotting analysis of the expression of the indicated proteins involved in cGAS-SITNG pathway of cortical tissues in 6-month-old *Cgas*^+/+^;5×FAD and *Cgas*^−/−^;5×FAD mice. *n* = 3. **k**, Quantification of the expression of p-STING), p-TBK1, p-p65, and p-IRF3 relative to β-actin in **j**. Mean ± SD; *n* = 3; *, *p* < 0.05, ***, *p* < 0.001, Student’s *t*-test. **l**, Transcriptional analysis of a panel of A1 astrocyte-inducing, disease-associated microglial markers, and interferon (IFN)-stimulated genes in hippocampal tissues of 6-month-old *Cgas*^+/+^;5×FAD and *Cgas*^−/−^;5×FAD mice compared to WT mice. *n* = 4.

Aβ deposits activates glia cells which, in turn, can aggravate Aβ pathology^34^. Immunostaining with microglial marker Iba1 showed the reduced microglial activation in DG (Fig. 2h and Extended Data Fig. 2f) tissues of *Cgas*^−/−^;5×FAD mice compared to their *Cgas*^+/+^;5×FAD littermates. In addition, *Cgas* deficiency restored synaptic function, at least partially as shown by higher levels of synaptic proteins PSD95 and Synaptophysin in cortical tissues of *Cgas*^−/−^;5×FAD mice (Extended Data Fig. 2g, h). To uncover the effect of the cGAS-STING pathway on neuroinflammatory responses in 5×FAD mice, we examined the protein components in the cGAS-STING pathway and found the decreased levels of p-STING, p-TBK1, p-p65 and p-IRF3 in cortical tissues of *Cgas*^−/−^;5×FAD mice (Fig. 2j, k). Furthermore, we performed transcriptional analysis of a panel of genes^35^ that are highly expressed in AD-associated microglia^36^. The results showed the decreased expression of these genes, such as inflammatory cytokines including *Tnfa* and *Il1a*, complement factors including *C1q, C3* and *C4a*, and IFN-stimulated genes such as *Ifit1* and *Oas1*, in hippocampal tissues of *Cgas*^−/−^;5×FAD mice relative to their *Cgas*^+/+^;5×FAD littermates (Fig. 2l). Thus, these results suggest a previously unknown role of the cGAS-STING pathway in AD pathogenesis via 5×FAD mice.

### Inhibition of neurotoxic A1 astrocytes by cGAS deficient microglia

To dissect the cGAS-STING pathway in different brain cell types, we cultured primary microglia, neurons, and astrocytes, which were treated with oligomeric human Aβ42 peptides. As a result, we found the consistent increase of cmtDNA levels in these cells compared to untreated (mock) group, with the relative higher cmtDNA levels in neurons (Extended Data Fig. 3a-f). Accordingly, we observed the increased level of 2′3′-cGAMP, the direct product of cGAS sensing dsDNA, from these primary cells (Extended Data Fig. 3g-i). In examining the consequence of the activation of the cGAS-STING pathway, we found the marked increase of IFN-β level in the supernatants of primary microglia but not in primary neurons or astrocytes upon the treatment of oligomeric Aβ42 peptides (Extended Data Fig. 3j-l). These results suggest that Aβ oligomers can induce broad activation of cGAS in multiple neural cells, yet relatively specific STING-IFN response in microglia.

We next validated cGAS-STING activation in primary microglia cultures after oligomeric Aβ42 treatment (Fig. 3a). Immunoblotting with the cell lysates showed the increased p-TBK1, p-p65 and p-IRF3 in Aβ42-treated microglia relative to mock (Fig. 3b, c). ELISA assays revealed higher levels of inflammatory cytokines including IL-6 (Fig. 3d), TNF-α (Fig. 3e), and IL-1α (Fig. 3f) in Aβ42-treated *Cgas*^+/+^ microglia than mock, and, however, *Cgas* deficiency caused substantial decreases of these inflammatory cytokines in the same condition. We next examined the impact of *Cgas* deficiency in different neural cells on Aβ42-induced neuronal damage. Firstly, *Cgas*^+/+^ or *Cgas*^−/−^ mouse primary neurons were treated with oligomeric Aβ42 and showed comparable levels of neuronal damage (Extended Data Fig. 4a-c). Next, we applied astrocyte-conditioned medium (ACM) or microglia-conditioned medium (MCM) collected from *Cgas*^+/+^ or *Cgas*^−/−^astrocytes or microglia, respectively, to mouse primary neurons cultures challenged with oligomeric Aβ42 peptides, and found neither *Cgas*^−/−^ ACM nor *Cgas*^−/−^ MCM had differential effect on neuronal survival when compared with *Cgas*^+/+^ ones (Extended Data Fig. 4d-i), suggesting the phenotypes observed *in vivo* are likely due to more complex intercellular crosstalk.

**Fig. 3:**
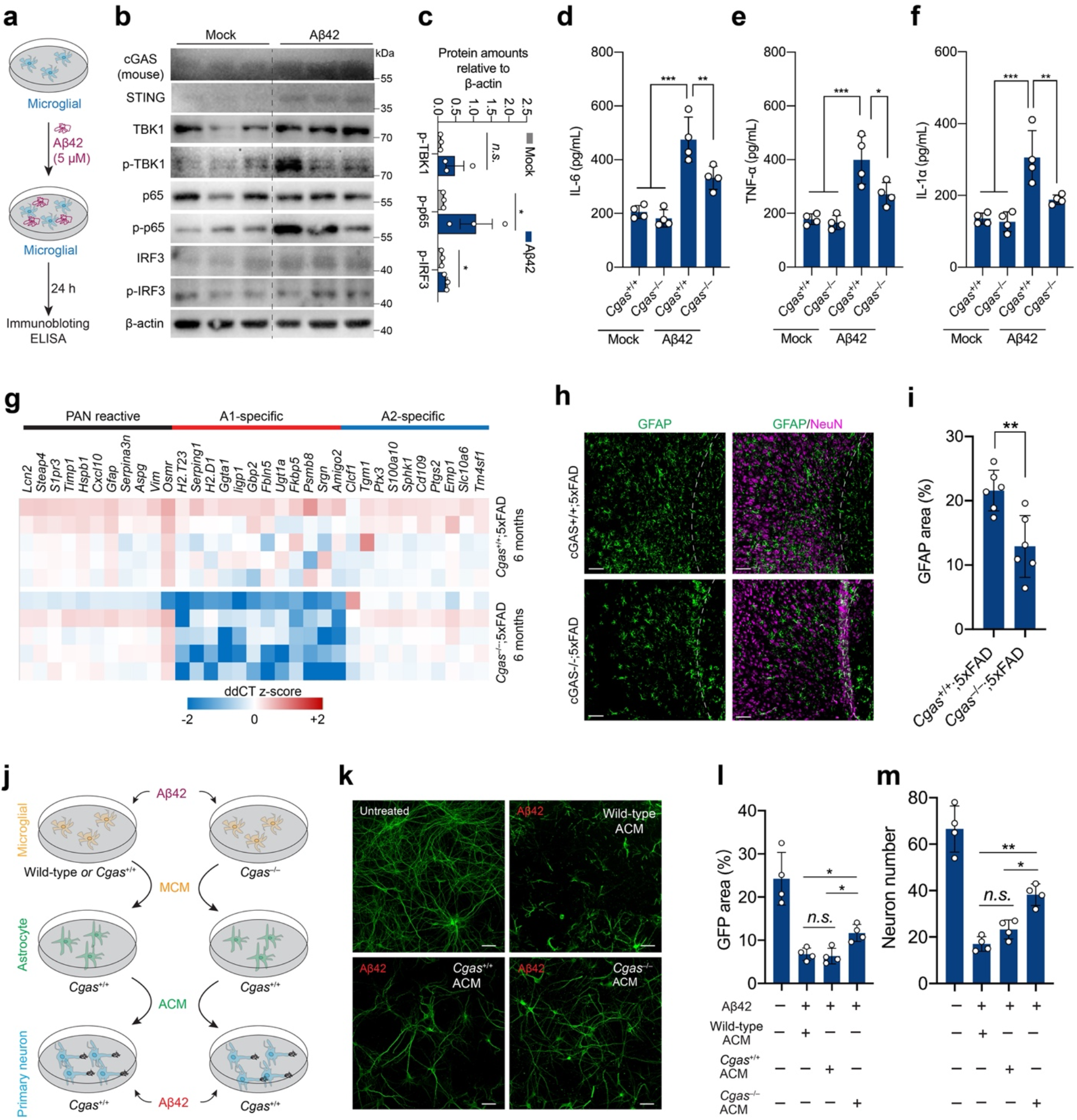
The suppression of neurotoxic A1 astrocytes by cGAS deficient microglia is neuroprotective from Aβ toxicity. **a**, Schematic diagram showing primary cultured microglia treated with oligomeric Aβ42 (5 μM) for 24 h, followed by immunoblotting and ELISA analysis. **b**, Immunoblotting analysis of the expression of the indicated proteins involved in cGAS-SITNG pathway of primary WT microglia 24 h after the Aβ treatment. The non-treatment of the Aβ was used as control. *n* = 3. **c**, Quantification of the expression of p-TBK1), p-p65, and p-IRF3 relative to β-actin in **b**. Mean ± SD; *n* = 3; *n*.*s*., not significant; *, *p* < 0.05, Student’s *t*-test. **d-f**, ELISA analysis for IL-6 (**d**), TNF-α (**e**) and IL-1α (**f**) in the supernatants of primary *Cgas* ^+/+^ or *Cgas* ^−/−^ microglia 24 h after Aβ42 treatment. The non-treatment of Aβ42 was used as control. Mean ± SD; *n* = 4; *, *p* < 0.05, **, *p* < 0.01, ***, *p* < 0.001, one-way ANOVA with Bonferroni’s *post hoc* test. **g**, Transcriptional analysis for activated astrocytic genes in 6-month-old *Cgas*^+/+^;5×FAD (*n* = 5) and *Cgas*^−/−^;5×FAD (*n* = 5) mice. A1-specific: genes activated only by LPS; A2-specific: genes activated only by ischaemia; Pan-reactive: genes activated by either LPS or ischaemia. **h**, GFAP^+^ astrocytic staining of cortical region in 6-month-old *Cgas*^+/+^;5×FAD and *Cgas*^−/−^;5×FAD mice. Scale bar, 50 μm. **i**, Quantification of percentage of GFAP^+^ area in **h**. Mean ± SD; *n* = 6; **, *p* < 0.01, Student’s *t*-test. **j**, Schematic diagram showing WT, *Cgas*^+/+^, and *Cgas*^−/−^ primary microglia treated with oligomeric Aβ42 (5 μM) for 24 h, the resulted microglia conditioned medium (WT MCM, *Cgas*^+/+^ MCM, and *Cgas*^−/−^ MCM, respectively) individually added to *Cgas*^+/+^ primary astrocytes for inoculation for 24 h, and the concentrated astrocyte conditioned medium (WT ACM, *Cgas*^+/+^ ACM, and *Cgas*^−/−^ ACM, respectively)) individually inoculated with *Cgas*^+/+^ primary neurons treated in parallel with oligomeric Aβ (750 nM) for 24 h. Primary neurons were pre-treated with adenovirus-associated virus (AAV) expressing a GFP with CaMKII promotor. **k**, Fluorescent signals in primary neuron cultures treated with oligomeric Aβ, together with WT, *Cgas*^+/+^, or *Cgas*^−/−^ACM. Scale bar, 50 μm. **l, m**, Percentage of GFP-labeled primary neurons (**l**) and numbers of primary neurons (**m**) in **k**. Mean ± SD; *n* = 4; *n*.*s*., not significant; *, *p* < 0.05, **, *p* < 0.01, one-way ANOVA with Bonferroni’s *post hoc* test.

Notably, TNF-α and IL-1α have been considered as the critical inflammatory factors secreted from activated microglia to convert astrocytes to neurotoxic A1 astrocytic phenotype^37,38^. Thus, we are stimulated to examine the effect of the cGAS-STING pathway on astrocytic subtyping *in vivo* by purifying ACSA-2^+^ astrocytes from brains of 6-month-old mice (Extended Data Fig. 5a-c). Transcriptional analysis of these purified astrocytes showed overall no significant difference regarding the expression of pan-reactive genes, A1 specific genes and A2 specific genes between *Cgas*^+/+^ and *Cgas*^−/−^ mice (Extended Data Fig. 5d). However, the results revealed a modest decrease of the expression of some pan-reactive genes like *Gfap*, a significantly reduced expression of neurotoxic A1 specific genes such as *H2*.*T23* and *Serping1*, and an overall unchanged expression of neuroprotective A2 specific genes like *Clcf1* and *Tgm1* in the purified astrocytes from *Cgas*^−/−^;5×FAD mice compared to their *Cgas*^+/+^;5×FAD littermates (Fig. 3g). Consistently, immunostaining revealed the reduction of GFAP^+^ astrocytic activation in cortical (Fig. 3h, i) and DG (Extended Data Fig. 5e, f) tissues of *Cgas*^−/−^;5×FAD mice relative to their *Cgas*^+/+^;5×FAD littermates.

As *Cgas* deletion reduced the expression of the microglia-derived factors that induce A1 astrocytes, we next examined the effect of the cGAS-STING pathway-regulated astrocytic subtyping on neuronal survival from Aβ toxicity *in vitro*. To examine the potential effect of A1 astrocytes on neuronal damage, *Cgas*^+/+^ or *Cgas*^−/−^ MCM after oligomeric Aβ42 treatment was individually added to WT astrocyte cultures for 24 h, and, subsequently, the resulted *Cgas*^+/+^ ACM, or *Cgas*^−/−^ ACM was applied to WT primary neurons treated in parallel with oligomeric Aβ42 (Fig. 3j). As a result, *Cgas*^+/+^ ACM failed to improve oligomeric Aβ42-induced neuronal cell death in primary neurons. However, neuronal cell death induced by oligomeric Aβ42 was significantly reduced after *Cgas*^−/−^ ACM treatment (Fig. 3k), as quantified by GFP signal (Fig. 3l) and neuron numbers (Fig. 3m). These results demonstrate that cGAS-STING regulates neuroinflammatory response and neuronal health through a tri-cellular signaling pathway.

### Pharmacological inhibition of the cGAS-STING pathway ameliorates Aβ pathology in 5×FAD mice

To explore the potential of treating AD by modulating the activation of the cGAS-STING pathway, we first performed *in vitro* experiments in the tri-cellular setting (Fig. 4a). As a result, cGAS inhibitor RU.521 ACM and STING inhibitor H-151 ACM provided strong neuroprotection from oligomeric Aβ42-induced neuronal toxicity (Fig. 4b) as similarly quantified by GFP signal (Fig. 4c) and neuron numbers (Fig. 4d). However, neither RU.521 MCM nor H-151 MCM had significant protective effect on neuronal cells alone when challenged with Aβ42 toxicity (Extended Data Fig. 6a-c). Next, we administrated STING inhibitor H-151 in 3-month-old 5×FAD mice to pharmacologically inhibit STING-IFN activation as previously reported in other neurological disorder^21^ (Fig. 4e). H-151 treatment suppressed the activation of the cGAS-STING pathway in the brains, as indicated by the reduced levels of p-TBK1, p-p65 and p-IRF3 proteins (Extended Data Fig. 7a, b). Administration of H-151 reduced the levels of Aβ42 in TBS and guanidine fractions, but not Aβ40 in both fractions, in cortical tissues of 5-month-old 5×FAD mice compared to vehicle treatment (Fig. 4f-i). Furthermore, immunostaining with thioflavin S showed the reduction of Aβ cores in DG and cortical tissues of H-151-treated 5×FAD mice relative to vehicle group (Fig. 4j-n). Additional immunostaining revealed the decrease of Aβ deposits (Fig. 4o, p) and the reduction of Iba1^+^ microglia (Fig. 4o, q) and GFAP^+^ astrocytes (Fig. 4o, r) in DG of H-151-treated 5×FAD mice compared to vehicle control. Consistently, H-151 treatment markedly inhibited the expression of a panel of neuroinflammatory genes including *Tnfa, C1q, Il1a*, and *C3* in hippocampus of 5×FAD mice (Extended Data Fig. 7c-f). Nevertheless, H-151 administration did not affect the number of NeuN^+^ neurons in DG of 5×FAD mice (Extended Data Fig. 8a-c), as neuronal loss usually occurs after 6 months in this mouse model. Additionally, we observed that H-151 treatment enhanced microglial phagocytosis activity as shown by the relative increase of CD68/Aβ ratio compared to vehicle control (Extended Data Fig. 8d, e). Therefore, these results revealed that H-151-mediated pharmacological inhibition of the cGAS-STING pathway ameliorated AD pathogenesis, suggesting a therapeutic that blocks the activity of the cGAS-STING pathway might effectively interfere with the progression of AD.

**Fig. 4:**
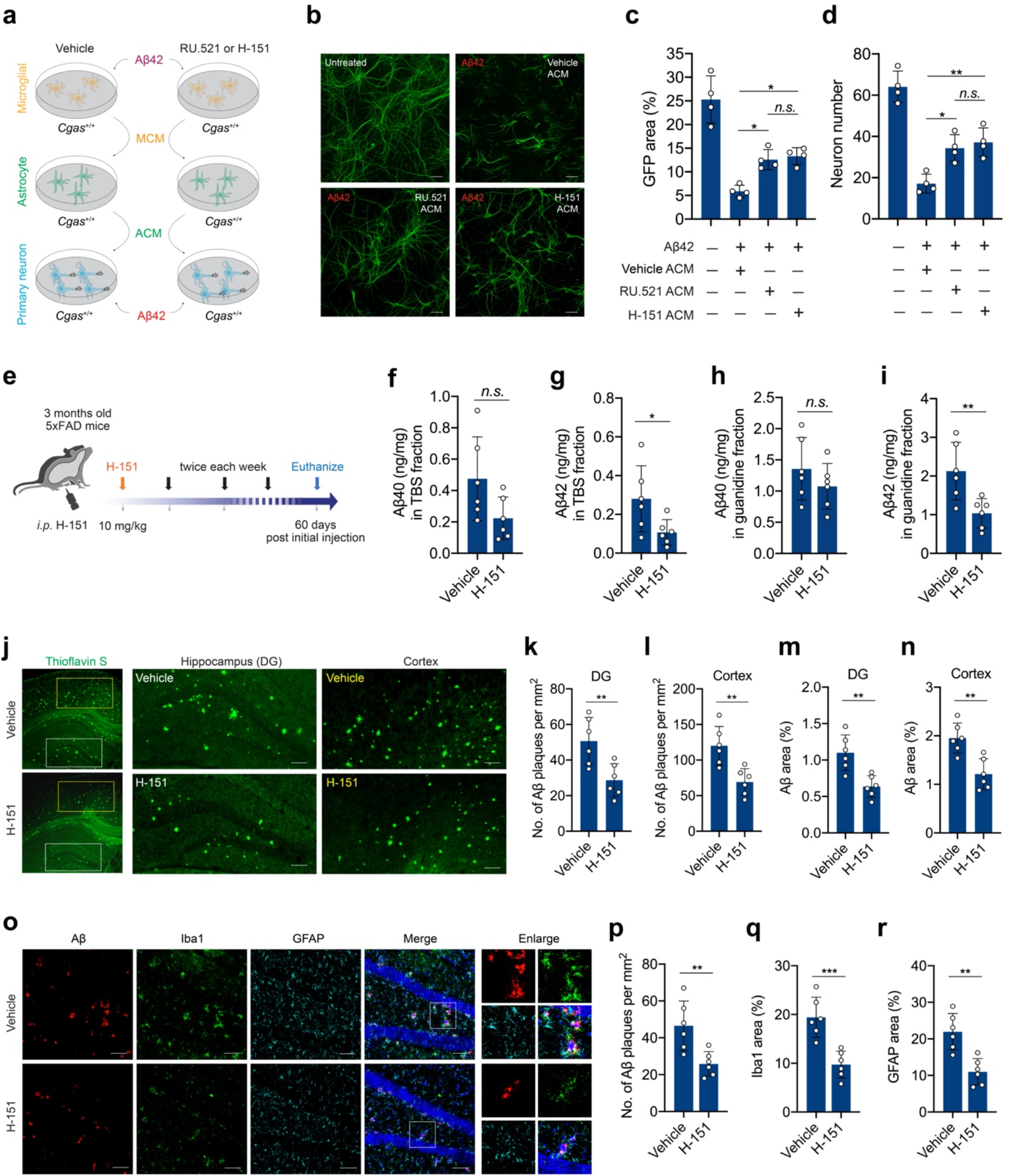
Administration of STING inhibitor H-151 ameliorates Aβ pathology and neuroinflammation in 5×FAD mice. **a**, Schematic diagram showing *Cgas*^+/+^ primary microglia treated with oligomeric Aβ42 (5 μM) for 24 h, together with cGAS inhibitor RU.521 (50 μM) or STING inhibitor H-151 (15 μM), the resulted microglia conditioned medium (MCM) added to *Cgas*^+/+^ primary astrocytes for inoculation for 24 h, and the concentrated astrocyte conditioned medium (ACM) inoculated with cGAS^+/+^ primary neurons treated in parallel with oligomeric Aβ (750 nM) for 24 h. Primary neurons were pre-treated with adenovirus-associated virus (AAV) expression a GFP with CaMKII promotor. **b**, Fluorescent signals in primary neuron cultures treated with oligomeric Aβ42, together with the RU.521 or H-151 ACM. Scale bar, 50 μm. **c, d**, Percentage of GFP-labeled primary neurons (**c**) and numbers of primary neurons (**d**) in **b**. Mean ± SD; *n* = 4; *n*.*s*., not significant; *, *p* < 0.05, **, *p* < 0.01, one-way ANOVA with Bonferroni’s *post hoc* test. **e**, Schematic diagram of STING inhibitor H-151 treatment on 3-month-old 5×FAD mice (see Methods). **f-i**, Quantification of the indicated Aβ40 (**f**) and Aβ42 (**g**) in TBS fraction, and Aβ40 (**h**) and Aβ42 (**i**) in guanidine fraction of cortical tissues in H-151-treated 5-month-old 5×FAD mice by ELISA. Mean ± SD; *n* = 6; *n*.*s*., not significant; *, *p* < 0.05, **, *p* < 0.01, Student’s *t*-test. **j**, Thioflavin S staining of DG and cortical tissues in H-151-treated 5-month-old 5×FAD mice. Scale bar, 50 μm. **k, l**, Quantification of Thioflavin S-labeled amyloid core numbers per mm^2^ in DG (**k**) and cortical (**l**) regions in **j**. Mean ± SD; *n* = 6; **, *p* < 0.01, Student’s *t*-test. **m, n**, Quantification of percentage of Thioflavin S-labeled Aβ plaque area in DG (**m**) and cortical (**n**) regions in **j**. Mean ± SD; *n* = 6; **, *p* < 0.01, Student’s *t*-test. **o**, Immunostaining of Aβ (6E10), Iba1^+^ microglial and GFAP^+^ astrocytes of DG region in H-151-treated 5-month-old 5×FAD mice. Scale bar, 50 μm. **p-r**, Quantification of Aβ plaque numbers per mm^2^ (**p**) and percentage of Iba1^+^ (**q**) and GFAP^+^ (**r**) area in **o**. Mean ± SD; *n* = 6; **, *p* < 0.01, ***, *p* < 0.001, Student’s *t*-test.

## Discussion

Chronic activation of innate immunity is a fundamental trigger for neuroinflammation, which is critically biological process in the development and progression of AD. In the present study, we have identified cGAS-dsDNA binding in the cytosol of neural cells and, as a result, activation of the cGAS-STING pathway during AD and aging. We also provide evidence that the genetic deletion of *Cgas* gene remarkably ameliorates AD pathogenesis in 5×FAD mouse model, which is at least partially mediated by the suppression of neurotoxic A1 astrocytes. Therefore, our data support the hypothesis that the activation of the cGAS-STING pathway induced by cytoplasmic dsDNA, for example cmtDNA, accelerates AD progression. Inflammatory mediators that result from cGAS-STING activation are probably involved in the restriction of beneficial microglial clearance, conversion of neurotoxic A1 astrocytic phenotype, neural dysfunction, and cognitive impairment (Extended Data Fig. 9). This key role of the cGAS-STING pathway within innate immunity in the neurodegenerative progression suggests that a therapeutic that blocks the activity of cGAS-STING signaling, or the downstream inflammatory mediators, might effectively interfere with the progression of AD.

Neuroinflammation is increasingly considered as one of the most harmfully pathological processes in the development and progression of AD^39-42^, and closely interacted with other AD pathological factors such as Aβ and tau^43^. Neuroinflammation is usually caused by an uncontrolled or excessive innate immune response, most commonly from microbial infections. However, molecular evidence directly linking innate immunity with neurodegenerative diseases remains largely underexplored. Our present study has provided evidence that the cGAS-STING pathway is evidently activated during AD and aging, and consequently contributes to AD pathogenesis, as shown by cognitive evaluation, Aβ pathology, and neuroinflammatory response via *Cgas*^−/−^;5×FAD mice. Furthermore, the treatment of STING inhibitor H-151 can effectively suppress the activation of the cGAS-STING pathway *in vivo*, as shown by the decrease of the pathway protein phosphorylation and inflammatory mediators, and attenuate AD pathogenesis at least in 5×FAD mouse model. A recent study^24^ suggests the involvement of the cGAS-STING pathway in tau pathology. Combining with our present investigation on amyloid pathology, all these outcomes together demonstrate that cGAS-STING as a critical innate immune controller plays an important role in AD pathogenesis and is a promising immune target for AD intervention. Notably, other critical innate immunity sensors including NLRP3 inflammasome have been shown to be involved in AD pathogenesis^44,45^. Therefore, all these critical molecular links between innate immunity and AD may represent a new hypothesis.

Within brain cells, neurons are susceptible to the toxicity of the neurodegenerative proteins like Aβ and hyperphosphorylated tau, which cause a harmful stress to adversely affect multiple cellular biological functions, including mitochondrial homeostasis and DNA stability^26,27^. Consistently, we found the cytosolic cGAS-dsDNA binding in human and mouse AD and aged mice. However, the activation of the SITNG-IFN cascade following cGAS sensing cytosolic dsDNA was present mainly in microglia but not neurons and astrocytes as shown in our present and previous studies^29,30^. This is triggering a question of how the signal of cGAS sensing dsDNA within neural cells is transduced to the activation of microglial STING-IFN cascade. One of the potential clues has been reported that 2’3’-cGAMP produced by cGAS upon dsDNA sensing can be transported between neighboring cancer cells via specific transporters^46,47^ and gap junction^48^, suggesting that 2’3’-cGAMP produced in neural cells could be somehow converged into microglia and subsequently induce the activation of the STING-IFN cascade in microglia, thereby stimulating neuroinflammatory responses. Thus, understanding the mechanism underlying cGAS-STING signal transduction between neural cells would be helpful to look for approaches to modulate such process, in parallel to block the activity of the cGAS-STING pathway, thereby jointly interfering with the progression of AD. It should be noted that, the cGAS-STING pathway-targeting therapies, besides H-151 used in the present study, deserve to be further examined for AD intervention.

In conclusion, we have identified the cGAS-STING pathway as a critical molecular link between innate immunity and AD and uncovered the complicated interactions between microglia, astrocytes, and neurons among neuroinflammatory microenvironment. Future studies are warranted to further explore how the neurodegenerative changes are elaborately modulated by innate or adaptive immunity, and if these harmful effects on neuronal function can be partially improved or even reversed.

## Methods

### Human postmortem brain samples

Frozen and paraffin-embedded human postmortem brain samples from patients with AD and healthy controls (age-matched and died of certain cause unrelated to dementia) were acquired from the Chinese Brain Bank Center (CBBC), Wuhan, China. The detailed demographic information is present in Supplementary Table 1. The experimental procedures with human postmortem brain tissues were reviewed and approved by the Ethics Board at Kunming Institute of Zoology (KIZ), Chinese Academy of Sciences (approval no. KIZRKX-2022-003). The brain samples were quickly frozen and used for immunoblotting analysis. In addition, brain tissues were fixed and embedded into paraffin and five-micrometer thick consecutive sections on silane-coated slides were used for immunohistochemistry.

### Mice

*Cgas*^−/−^ mice (B6(C)-*Cgas*^tm1d(EUCOMM)Hmgu^/J) (Stock no: 026554) under the control of cytomegalovirus (CMV) enhancer fused to the chicken beta-actin promoter (CAG) promotor were purchased from the Jackson Laboratory. 5×FAD mice (Stock no: 008730) and C57BL/6J mice (Stock no: 000664) were purchased from the Jackson Laboratory. *Cgas*^−/−^ mice were mated with 5×FAD mice to produce heterozygous *Cgas*^*+*/−^;5×FAD, and *Cgas*^*+*/−^;5×FAD mice were used to generate *Cgas*^−/−^;5×FAD and *Cgas*^+/+^;5×FAD littermates. As the difference in Aβ pathology and neuroinflammation between male and female 5×FAD mice was reported^49^, male mice were used throughout the study except additional clarification. 3-month-old 5×FAD mice were intraperitoneally injected with STING inhibitor H-151 (10 mg/kg body weight) (S6652, Selleck) twice each week. Dimethylsulfoxide (DMSO) that dissolved H-151 was used as vehicle control with the same volume for intraperitoneal injection for 3-month-old 5×FAD mice. The effects of H-151 on neurological outcomes in the mice were assessed at 2 months after initial H-151 administration. Animals were maintained on a 12-h light/dark cycle, with free access to food and water. The Institutional Animal Care and Use Committee of the KIZ (approval no. IACUC-RE-2022-04-004) approved all experimental procedures and protocols.

### Behavioral test

The behavioral tests were performed as previously described^50^. To evaluate nest construction behavior, mice were individually placed in their home cages with a pre-weighed nestlet 1 h before the dark phase. The nests were assessed in the next morning and scored 1∼5^51^. The unused nestlet was weighed to count the percentage of nestlet used. To assess burrowing behavior, mice were individually placed in cages that were placed with a 200-mm long and 70-mm diameter burrow. The burrow was pre-filled with 200 g of mouse food pellets. Mice were placed in the cages to burrow for 3 h. The remaining food pellets inside the burrow were weighed to count the amount burrowed.

### Isolation of astrocytes from adult mouse brains

Adult mice were anesthetized and perfused transcardially with ice-cold saline. Whole brain without cerebellum was operated to remove meninges in prechilled phosphate-buffered saline (PBS). Brain tissues were mechanically dissociated and prepared for sing-cell suspension by using the Adult Brain Dissociation kit (130-107-677, Meltenyi Biotec) accordingly to the manufacture’s instruction. Myelin in the suspension was removed using the cell debris removal buffers. The obtained single-cell pellets were resuspended with PBS containing 1% bovine serum albumin (BSA). Astrocytes were isolated using anti-ACSA-2 MicroBeads (130-097-678, Meltenyi Biotec) by using the MACS multi-stand separator accordingly to the manufacture’s instruction.

### Isolation and culture of primary neuron, microglia and astrocyte

Primary hippocampal neurons were isolated from embryonic day 18.5 pups and cultured in Neurobasal medium (Gibco) supplemented with 1% B-27, 0.5 mM L-glutamine, 100 U/mL penicillin-streptomycin on tissue-culture plates coated with poly-L-lysine. The medium was changed to maintain neuron cultures every 3 days as previously described^52^. The isolation and culture of primary microglia and astrocyte were conducted as described previously^38^. Whole mouse brains without cerebellum from postnatal day 1 (P1) pups were used to remove the meninges. The remained brain tissues were then washed in Dulbecco’s modified Eagle’s medium (DMEM)/F12 supplemented with 10% heat-inactivated FBS, 1% penicillin and streptomycin, 2 mM L-glutamine, 100 μM non-essential amino acids (NEAA), and 2 mM sodium pyruvate (DMEM/F12 complete medium) three times. The brains were mechanically dissociated, followed by the incubation of 0.25% trypsin-EDTA for 10 min. The DMEM/F12 complete medium was added to stop the trypsinization and a single-cell suspension was prepared by trituration. A 100-μm nylon mesh was used to remove cell debris and aggregates within the single-cell suspension. The obtained single cell was cultured in T75 flasks for 8∼10 days, with the complete medium replacement on day 4. The glial cells were separated into astrocytic and microglial fractions by using the anti-ACSA-2 MicroBeads (130-097-678, Meltenyi Biotec). The obtained astrocytes and microglia were cultured separately. The conditioned medium from the primary activated microglia (MCM) treated with 5 μM oligomeric Aβ (4090148.1000, Bachem) were collected and added to primary astrocytic cultures for 24 h. The conditioned medium from the treated astrocytes (ACM) were collected with protease inhibitor (K1007, Apexbio), followed by the concentration with Amicon Ultra-15 centrifugal filter unit (10 kDa cutoff) (Millipore). 15 μg/mL of total concentrated proteins determined by a BCA protein assay kit (P0010S, Beyotime) were added to mouse primary neuron cultures, pre-treated with adenovirus-associated virus (AAV) expressing a GFP with CaMKII promotor, for neuronal survival assay.

### Quantitative real-time PCR (qRT-PCR)

Total RNA was extracted using a TRIzol reagent (15596018, ThermoFisher Scientific) according to the manufacturer’s instructions. RNase-free DNase (M6101, Promega) was used for DNase digestion during RNA purification. RNA quantity and quality were assessed by Nanodrop for RNA isolated from tissues. cDNA was generated from 1 μg total RNA per sample using the M-MLV Reverse Transcriptase kit (M1705, Promega). To detect RNA transcript of targeted genes, the target transcripts were determined by qPCR using the PereCTa SYBR Green SuperMix (95055-500, Quanta Biosciences) on CFX96 real-time PCR system (Bio-Rad). The primer sequences are listed in Supplementary Table 2.

### Immunofluorescence

Immunofluorescence was performed as described in our previous study^53^. Briefly, brain slides between bregma -1.8 mm and -2.2 mm were prepared via frozen sections for each animal. For antigen retrieval, slides were immersed in a quick antigen retrieval solution (P0090, Beyotime). Then slides were washed with 1 × PBS (pH 7.4), and blocked with 5% BSA in 1 × PBST (0.3% Triton-X 100 in PBS) at 37°C for 60 min. The primary antibodies used are listed in Supplementary Table 3. The primary antibodies above were diluted in 3% donkey serum in 1 × PBST (0.2% Triton-X 100) and incubated overnight at 4°C. Slides were then washed, and immunoreactivity was detected using Donkey anti-Rabbit IgG Highly Cross-Adsorbed Secondary Antibody, Alexa Fluor Plus 488, Donkey anti-Mouse IgG Highly Cross-Adsorbed Secondary Antibody, Alexa Fluor Plus 555, Donkey anti-Rat IgG Highly Cross-Adsorbed Secondary Antibody, Alexa Fluor Plus 647 (1:500; ThermoFisher Scientific) for 1 h at room temperature. Slides were counterstained with 5 μg/mL 4’,6-diamidino-2-phenylindole (DAPI; ThermoFisher Scientific) for 10 min at room temperature and washed with 1 × PBST (0.2% Triton-X 100) three times. Slides were visualized using the Nikon A1R confocal microscope and then analyzed by ImageJ software.

### Immunoblotting

Immunoblotting was performed as previously described^54^. In brief, protein samples were separated by SDS-PAGE and transferred to PVDF membrane by semi-dry transfer at 25 V for 30 min. The membrane was blocked in 5% skim milk in 1 × PBST for 1 h and incubated overnight with commercial primary antibody in 5% BSA at 4°C. The membrane was incubated with anti-mouse or anti-rabbit HRP-conjugated secondary antibodies in 5% milk and bands were developed with Chemi-Doc XRS imaging (Bio-Rad). The primary antibodies used are listed in Supplementary Table 3.

### Flow Cytometry

Cell samples were fluorescently stained with Anti-ACSA-2-PE (clone: IH3-18A3) (130-102-365, Miltenyi Biotec), examined by a BD FACSCanto II flow cytometer, and analyzed with FlowJo software.

### Enzyme-linked immunosorbent assay (ELISA)

The secreted cytokines from primary neural cells after oligomeric Aβ42 treatment were quantified by mouse cytokine ELISA kits, including mouse IL-6 ELISA kit (E-EL-M0044c, Elabscience), mouse TNF-α ELISA kit (E-MSEL-M0002, Elabscience), mouse IL-1α ELISA kit (E-EL-M3059, Elabscience), and moue IFN-β ELISA kit (42410-1, PBL Assay Science) accordingly to the manufacture’s instruction.

Level of 2′3′-cGAMP within neural cell lysates after oligomeric Aβ42 treatment was measured by 2′3′-cGAMP competitive ELISA kit (501700, Cayman Chemical) accordingly to the manufacture’s instruction.

Plaque-related Aβ were extracted using the protocols previously described with minor modification^50^. Briefly, frozen cortical brain tissues were weighed, added with freshly prepared Tris-buffered saline (TBS) (20 mM Tris-HCL, 150 mM NaCl, PH 7.4), and homogenized on a mechanical homogenizer. The homogenate was centrifuged at 15,000 × g for 30 min at 4°C. The supernatants containing soluble Aβ were collected as TBS fraction for ELISA. The pellets were re-homogenized with an equal volume of TBS plus 5 M guanidine hydrochloride (pH 8.0) and incubated at room temperature. After re-centrifugation, the supernatants were collected as guanidine fraction for ELISA. The levels of Aβ40 and Aβ42 in mouse cortical tissues were determined by using Aβ40 kit (E-EL-H0542c, Elabscience) and Aβ42 kit (E-EL-H0543c, Elabscience), respectively, following the manufacturer’s protocols.

### Quantification of mtDNA release

Amount of mtDNA was quantified as previously described^55^. Briefly, isolated cells (1 × 10^6^) from adult mouse brains were resuspended in 170 μl of digitonin buffer containing 150 mM NaCl, 50 mM HEPES pH 7.4, and 25 μg/mL digitonin (HY-N4000, MedChemExpress). The homogenates were incubated on a rotator for 10 min at room temperature, followed by centrifugation at 15,000 × g for 25 min at 4°C. A 1:20 dilution of the supernatant (cmtDNA) was used for qPCR. The pellet was resuspended in 340 μl of lysis buffer containing 5 mM EDTA and proteinase K (HY-108717, MedChemExpress) and incubated at 55°C overnight. The digested pellet was diluted with water (1:20 to 1:100) and heated at 95°C for 20 min to inactivate proteinase K, and the sample was sued for qPCR with mtDNA specific primers listed in Supplementary Table 2. The cmtDNA in the supernatant was normalized to the total mtDNA in the pellet for each sample.

### Proximity ligation assay (PLA)

Binding of cGAS and dsDNA in human and mouse brain sections was determined by PLA kit (DUO92101, Sigma-Aldrich) accordingly to the manufacture’s instruction. The primary antibodies used in PLA include anti-dsDNA antibody (ab27156, Abcam) and anti-cGAS antibody (human-specific, 15102S, Cell Signaling Technology) or anti-cGAS antibody (mouse-specific, 31659S, Cell Signaling Technology). PLA signals were visualized using the Nikon A1R confocal microscope and indicated by PLA red dots per cell.

### Thioflavin S (TS) staining

Aβ plaques were labelled by TS staining on brain sections that were stained with 0.005% TS (T1892, Millipore) in 50% ethanol for 10 min. Then the sections were washed twice with 50% ethanol and with PBS three times. Brain sections were analyzed using the Nikon A1R or ZEISS LSM 880 confocal microscope.

## Statistical analysis

All appropriate data were analyzed using GraphPad Prism 8 (GraphPad Software Inc.). All hypothesis tests were performed as two-tailed tests. Specific statistical analysis methods were described in the related figure legends where results are presented. Values were considered statistically significant for *p* values < 0.05.

## Data availability

No sequencing data was produced in the present study.

## Declaration of interests

The authors declare no competing interests.

## Contributions

All authors read and approved the final version of the manuscript. JZ and ZZ conceived of the research and designed the study. JZ and ZZ wrote the manuscript. XX, GM, XL, and JBZ performed the experiments and discussed the data. All authors commented on the manuscript.

## Acknowledgements

We thank professor Ben Lu from the Central South University in China for providing additional *Cgas*^− /−^ mice and professor Yong-Han He from the Kunming Institute of Zoology, CAS for providing partial aged mice samples. The graphical diagram is produced by BioRender. This work was supported by Yunnan Fundamental Research Projects (grant No. 202201AW070020).

## Notes

### Competing Interest Statement

The authors have declared no competing interest.

